# Preparation of inactivated whole culture vaccine composed of *Pasteurella multocida, Avibacterium paragallinarum*, and *Ornithobacterium rhinotracheale* and evaluation of its protective efficacy in chickens

**DOI:** 10.1101/2022.11.25.517975

**Authors:** Sally Roshdy, Rafik Soliman, Manal A. Aly, Lamiaa Omar, Ahmed Samir, Heidy Abo-Elyazeed, Hassan Aboul-Ella

## Abstract

Poultry, mainly chickens and its white meat represents one of the main, nutritionally valuable, and affordable red meat replacer source of protein throughout the whole world with special reference to developing countries. A long list of microbial agents especially bacterial pathogens threat chickens production cycles. They constitute one of the major problems facing the rapidly expanding poultry industry and are responsible for considerable economic losses.). Fowl Cholera, Infectious Coryza, and Ornithobacterium Rhinotracheale (ORT) diseases were among the serious bacterial infections that affect respiratory tract of chickens with an global adverse effect on poultry production. A formalized whole culture vaccine of composed of *P. multocida* serotypes A5, A8, A9 and D2, *Avibacterium paragallinarum* serotype A and C and *Ornithobacterium rhinotracheale* serotype A was prepared. This polyvalent vaccine proved to be safe producing no adverse side effects when injected in chickens. The immunizing efficacy of this vaccine was evaluated in SPF chickens, which were immunized at 6 weeks of age. The protective efficacy of the vaccine was determined using challenge test. The developed vaccine was effective in protecting chickens against Fowl Cholera, Infectious Coryza and Ornithobacterium Infection (ORT diseases) in chickens against challenge with these pathogens. Vaccinated chickens challenged with virulent *Pasteurella Multocida* serotypes A5, A8, A9 and D2 showed protection rates of 86.6%, 93.3%, 93.3% and 93.3%, respectively, as compared with 100% mortality in the non-vaccinated control. Vaccinated chickens challenged with *Avibacterium paragallinarum* serotypes A and C showed protection rates of 86.6% and 93.3%, respectively. Also, the protection rate against challenge with virulent *Ornithobacterium rhinotracheale* serotype A reached to 96.6%.

## Introduction

Poultry represents an important and cheap source of protein throughout the world. The bacterial diseases of poultry, however, constitute one of the major problems facing the rapidly expanding poultry industry and are responsible for considerable economic losses **(Sharma, 1999; Bermudez and Stewart, 2008 and Cserep, 2008)**. Fowl Cholera, Infectious Coryza and ***Ornithobacterium Rhinotracheale*** (ORT) diseases were among the serious bacterial infections that affect respiratory tract of chickens **(OIE, 2008)**. Vaccines and vaccination strategies play an important role in controlling these diseases and minimizing the associated economic losses.

Fowl cholera is a contagious disease of domesticated and wild avian species caused by *Pasteurella multocida*. It occurs typically as a fulminating disease with massive bacteremia and high morbidity and mortality rates **(Rhoades and Rimler, 1990)**.

Infectious coryza is usually acute, sometimes chronic, highly infectious disease of chickens caused by the *Avibacterium paragallinarum* **(Blackall *et al*., 2005)** and characterized by catarrhal inflammation of the upper respiratory tract, especially nasal and sinus mucosa (**Paul McMullin, 2004)**.

*Ornithobacterium Rhinotracheale*, or ORT, is an acute highly contagious bacterial disease of birds, which is characterized by respiratory signs such as nasal discharge, sneezing, coughing, and sinusitis but in severe cases is followed by pneumonia, dyspnea, prostration, and mortality **(Van Empel and Hafez. 1999)**.

The effective prevention of these diseases depends upon the use of inactivated vaccines. The use of combined vaccines has the advantage of protection against more than one disease at the same time, beside reducing vaccination expenses, number of vaccinations performed and saving time. Therefore, the main objective of this study was to develop an inactivated combined adjuvanted, trivalent vaccine from the most virulent and locally prevalent serotypes of *Pasteurella multocida, Avibacterium paragallinarum* and *Ornithobacterium* rhinotracheale and to evaluate its immunizing and protective efficacy in chickens.

## Material and methods

### Experimental design (Fig.1)

In this study, 210 SPF chickens were used to evaluate the efficacy of the prepared trivalent vaccine. The chickens were divided into 3 groups:

**Figure (1):**
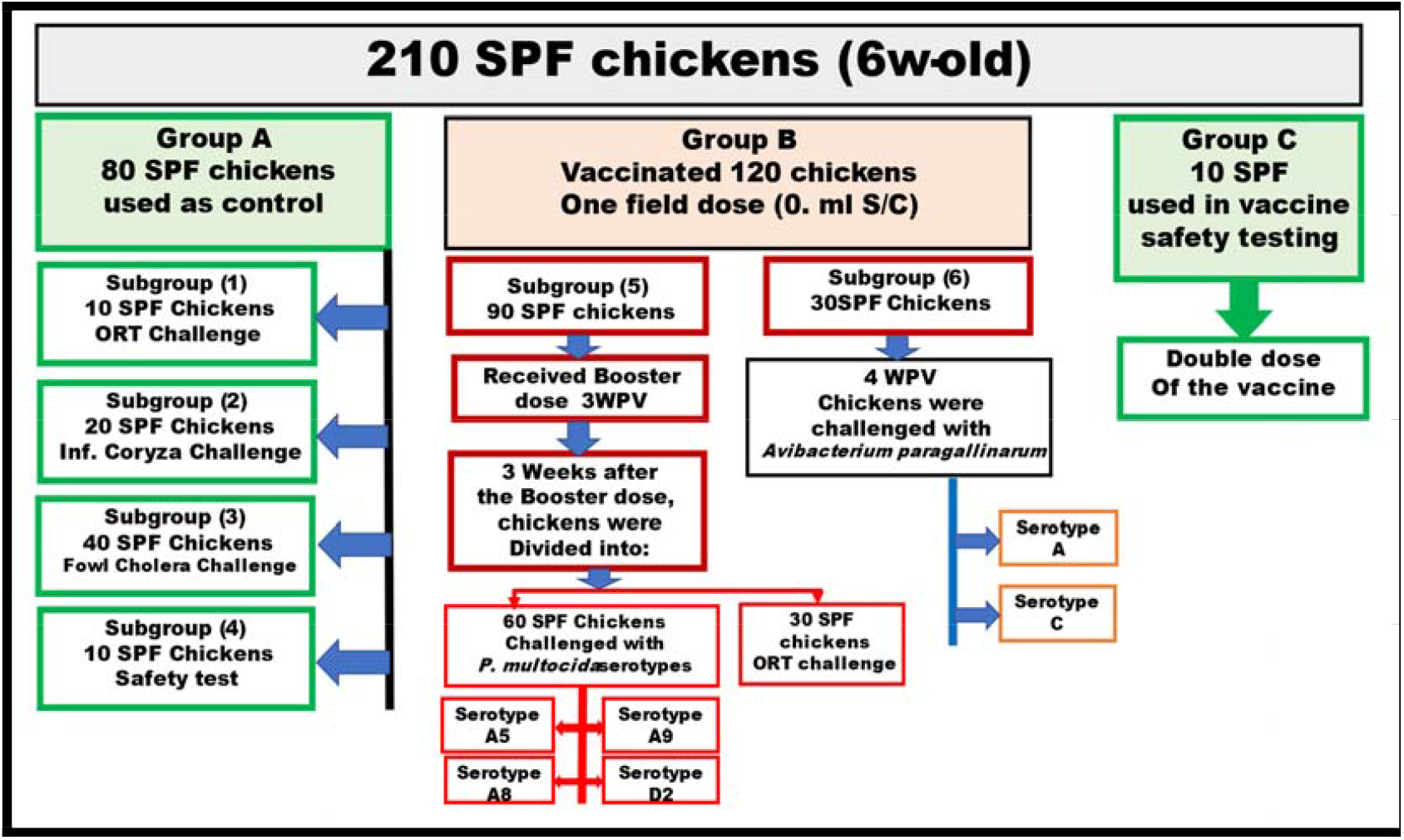
Flowchart to illustrate the overall SPF chickens subgrouping through the whole study

Group (A) was used as non-vaccinated control group and was divided into four subgroups which are 1, 2, 3, and 4 subgroups containing 10, 20, 40, and 10 SPF chickens, respectively. The subgroups 3, 2, and 1 were used for challenge test with fowl cholera, infectious coryza, and ORT, respectively. The SPF chickens in subgroup 4, however, was used for determination of safety. Group (B) was divided into two subgroups 5 and 6. The subgroup 5 contained 90 SPF chickens that were used for evaluation of the vaccine efficacy, and three weeks after immunization all the chickens in this subgroup received a booster dose. After further 3 weeks from the booster dose the immunized chickens were challenged with virulent fowl cholera serovars (15 chickens/each serovars) and with virulent *Ornithobacterium rhinotracheale* (30 chickens). While the subgroup 6 was used for challenge of immunized chickens at 4 weeks post immunization with *Avibacterium Paragallinarum* serotypes A and C (15 chickens/each serotype). Group (C) contained 10 SPF chickens that were injected with double the recommended dose of the vaccine for determination of its safety Figure (1).

### Vaccine seed bacterial strains: All the involved bacterial strains and serovars used for both vaccine preparation and further evaluation

*Pasteurella multocida* serotypes A5, A8, A9, and D2 obtained from the aerobic bacterial vaccines department, Veterinary Serum and Vaccine research institute (VSVRI), Abbasia, Cairo, Egypt. *Avibacterium paragallinarum* serotypes A and C that were supplied from Intervet International B.V. Boxmeer, Holland. *Ornithobacterium rhinotracheale* serotype A was obtained from the department of microbiology, faculty of veterinary medicine, Cairo University. It was used for the vaccine preparation and evaluation.

### Embryonated specific pathogen free chicken eggs (SPF-ECEs)

5-7 days old SPF-ECEs, were used for propagation of *Avibacterium paragallinarum*. These eggs were obtained from kom Oshem farm for SPF-ECEs, El-Fayuom, Egypt.

### Laboratory Animals

Experimental chickens: A total number of 210 chickens were used and obtained from (SPF) stocks from khom Oshem, El-Fayoum, Egypt. All involved chickens were housed in positive pressure stainless steel isolation cabinets at the central laboratory for evaluation of veterinary biologicals (CLEVB) with continuous light exposure and adliptum feeding strategy through the whole study period. Rabbits: Two months old Boskat rabbits with an average weight of 2.0 kg were used for the serial passage of *Pasteurella multocida* and preparation of hyper-immune sera.

□ **Formalin as inactivating agent (Hassan *et al*., 1992)** was obtained from BDH Limited Co., Poole, England. It was used in a form of formaldehyde solution 37% and was added to bacterial suspension for inactivation in 0.1% of final concentration.

□ **Adjuvants: Montanide** ^**TM**^ **ISA 70:** (SEPPIC-France) was obtained from **CLEVB**, and it used as W/O emulsion with a ratio 70:30.

□ **Bacterial culture media:**

#### 1. Brain Heart infusion broth

used for cultivation of *Pasteurella multocida* and *Ornithobacterium rhinotracheale*.

#### 2. Tryptose phosphate Broth (oxoid)

used for cultivation of *Avibacterium paragallinarum*.

## Methods

### Vaccine preparation

The bacterial components of the vaccine, namely, the *Pasteurella multocida* (serotypes, A5, A8, A9 and D2), *Avibacterium paragallinarum* serotypes A and C and *Ornithobacterium rhinotracheale* serotype A were grown separately in the culture media specific for each strain. The concentration of each *Pasteurella multocida* serotype (A5, A8, A9 and D2) culture was adjusted to 2×10^9^ CFU/ml as final concentration **(Singer and Malkinson, 1979)**, while the concentration of *Avibacterium paragallinarum* serotypes A and C was adjusted to 3×10^8^ CFU/ml as final concentrations **(Mohamed, 1996)**. The concentration of *Ornithobacterium rhinotracheale* serotype A culture was adjusted to 2×10^9^ **(Erganis *et al*., 2011)** as final concentrations.

### Vaccine inactivation with formalin (0.1%)

Formalin was added to inactivate each bacterial culture separately **(Hassan *et al*., 1992)**. The mixed antigens were absorbed with mineral oil and Montanid ISA70 at a ratio of 30:70, respectively **(Stone *et al*., 1978)**.

### Determination of the sterility and safety of the prepared vaccine

It was done according to the British veterinary codex 1970, European Pharmacopoeia 1997, Egyptian standard regulation for evaluation of veterinary biologics 2009, and OIE guidelines 2008 as follows; **Sterility test:** It was done by inoculating samples of the prepared vaccine on different bacterial and fungal media plates to determine if the prepared vaccine samples were free from bacterial, fungal, and Mycoplasma contamination or not.

#### Safety test

Ten healthy 6 weeks old SPF chickens were inoculated with twice the normal recommended dose of the prepared vaccine. The birds were observed for any possible local systemic adverse reactions for 21 days.

### Evaluation of the immunizing and protective efficacy of the prepared vaccine

Six weeks old SPF chickens were injected subcutaneously with 0.5ml of the prepared vaccine. As shown in the experimental design some chickens in group B were boosted at 3 weeks post vaccination with another field dose of the vaccine (**subgroup 5**). This subgroup was challenged at 3 weeks after the booster dose with the virulent serovars of *Pasteurella multocida* and *Ornithobacterium rhinotracheale* pathogens. While chickens in the **subgroup 6** remained without booster doses and were challenged at 4 weeks post immunization with virulent serovars of *Avibacterium paragallinarum* to determine the immunizing potential of the *Avibacterium paragallinarum* component of the tested trivalent vaccine.

The following bacterial culture doses were used for the challenge test of the vaccinated groups and non-vaccinated control chicken groups:

*Pasteurella multocida* group was challenged with a dose of 2×10^2^CFU/ml of for *Pasteurella multocida* virulent serovars via intramuscular route **(Egyptian standard regulation for evaluation of veterinary biologics. 2009)**.

In *Ornithobacterium rhinotracheale* group was challenged with 2 × 10^9^ CFU of ORT strain via spraying on the nose and eye **(Erganis *et al*., 2011)**.

In case of Infectious coryza group chickens were challenged virulent *Avibacterium paragallinarum* serotypes (A and C) with approximately 10^7^ CFU/0.5ml via infraorbital sinuses as well as via nostril route at a dose 0.2 ml **(Mohamed. 1996, Egyptian standard regulation for evaluation of veterinary biologics. 2009)**.

The challenged vaccinated and control groups chickens were put under observation for 14 days **(Erganis *et al*., 2011 and OIE, 2008]**. All challenged birds were observed daily after challenge and the morbidity and mortality rates were recorded for each group till the end of the observation period to measure the protection rate. Swabs from nostrils of challenged and control chickens were cultured for re-isolation of bacterial pathogens used in the challenge.

**Fig. 1: Experimental design of vaccination with the prepared vaccine and the challenge experiment**.

## Results

### 1 Safety and sterility of the prepared vaccine

The prepared trivalent vaccine was safe and free from bacterial and fungal contamination.

### 2 Results of challenge test

#### Protection efficacy of the prepared inactivated trivalent vaccine in chickens challenged with virulent *Pasteurella Multocida* serotypes A5, A8, A9 and D2

The data presented in **Table (1)**, indicated that chicken groups challenged with virulent *Pasteurella Multocida* serotypes A5, A8, A9 and D2 serovars showed protection rates of 86.6%, 93.3%, 93.3% and 93.3%, respectively. In the non-vaccinated control group 100 % of challenged chickens were died within of 3-4 days. Re-isolation of *Pasteurella Multocida* serotypes A5, 8, A9 and D2 was successful from control died chickens otherwise healthy vaccinated chickens showed negative results.

**Table (1):** Protection efficacy of the prepared vaccine in vaccinated chickens after challenge with virulent *Pasturella multocida* serotype (A5, A8, A9 and D2):

| Groups | *Pasteurella multocida* Serotypes | NO. of challenged birds | Mortality rate | Protection% |
| --- | --- | --- | --- | --- |
| Vaccinated | A5 | 15 | 2/15 | 86.6% |
|  | A8 | 15 | 1/15 | 93.3% |
|  | A9 | 15 | 1/15 | 93.3% |
|  | D2 | 15 | 1/15 | 93.3% |
| Control* | A5, A8,A9 and D2 | 10 | 10/10 | 0% |

Control\*:10 birds for each serotype.

#### Protection efficacy of the prepared trivalent inactivated vaccine in chickens challenged with *Avibacterium paragallinarum* serotypes A and C

In immunized chicken groups challenged with *Avibacterium paragallinarum* serotypes A and C, the protection rates reached to 86.6% and 93.3%, respectively. Chickens in the control non-vaccinated group that received the same challenge dose, 100 % of challenged chickens became diseased within a time of 3-4 days **(Table 2)**. Re-isolation of *Avibacterium paragallinarum* serotypes A and C was successful from control diseased chickens otherwise healthy vaccinated chickens showed negative results.

**Table (2).**
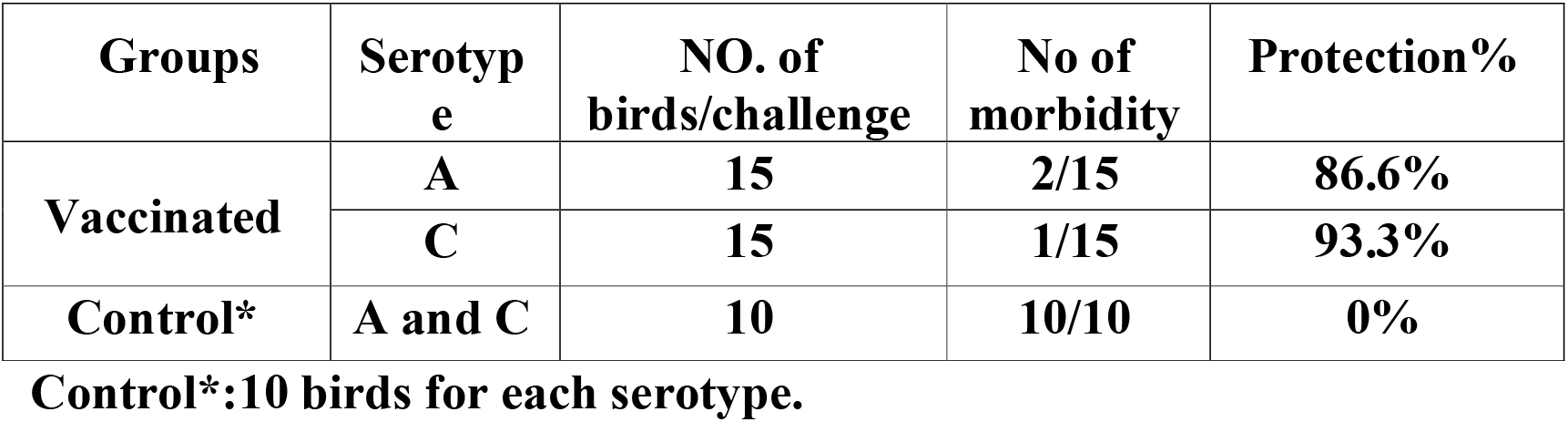
Protection rates in vaccinated chickens by challenged with *Avibacterium Paragallinarum* serotypes A and C.

#### Results of challenge test in immunized and non-vaccinated chickens challenged with *Ornithobacterium rhinotracheale* serotype(A)

As shown in **Table (3)**, vaccinated chicken groups challenged with *Ornithobacterium rhinotracheale* serotype A showed a protection rate of 96.6%. In the control non vaccinated group that received the same challenge dose, 90 % of challenged chickens were diseased. Re-isolation of *Ornithobacterium rhinotracheale* serotype A was successful from non-vaccinated control diseased chickens, otherwise healthy vaccinated chickens showed negative results.

**Table (3):** protection rates in chickens vaccinated with the prepared inactivated polyvalent vaccine after challenge with *Ornithobacterium rhinotracheale* serotype A.

| **Groups** | **Serotype** | **NO. of birds/challenge** | **No of morbidity** | **Protection%** |
| --- | --- | --- | --- | --- |
| **vaccinated** | **A** | **30** | **1/30** | **96.6%** |
| **Control** | **A** | **10** | **9/10** | **10%** |

## DISCUSSION

Fowl Cholera, Infectious coryza and ORT diseases stand clearly behind sever clinical respiratory distress, cough, sneezing and sinusitis in chickens. The major economic losses associated with these bacterial infection results from the rejection of carcasses for consumption, growth retardation and mortality **[OIE. 2008]**. The control of *Pasturella Multocida, Avibacterium paragallinarum* and ORT infections in chickens through vaccine have been described **[Gergis et al. 1992] [Kalayadari et al. 2004] [Jabbri and Moazani. 2005] [Erganis et al. 2010]**. Control of these diseases is very important and can be achieved by using vaccination strategies **[Gergis et al. 1992]**.

Preparation of combined inactivated vaccines is one of the tools for control of some bacterial respiratory diseases such as Fowl Cholera, Infectiuos Coryza and ORT diseases **[Bisschop et al. 2004] [Kalayadari et al. 2004] [Schuijfel et al. 2006][[Erganis et al. 2010]**. Inactivated vaccine adjuvanted with ISA70 montanid proved to induce long duration of immunity and high protective antibody titer **[Amal et al. 2001][Dungu et al. 2009][Ismail et al. 2013]**.

Safety and sterility testing of the prepared vaccine were carried out and proved that this vaccine had no adverse reactions and was free from bacterial and fungal contaminants. The use of formalin as inactivating agent together with the application of all safety procedure during the vaccine preparation stands behind this results.

Challenge under strictly condition may be also used to predict flock exposure response and can add considerable significance value obtained with the sera collected from the same chickens **[Stone. 1988]**.

Data presented in **Table (1)**, showed that all chickens immunized with the prepared vaccine were protected from infection with *Pasturella multocida* serotype A5, A8, A9 and D2 when challenged at the third week post injection of the booster dose with a protection rates of 86.6%, 93.3%, 93.3% & 93.3 %, respectively. This agreed with those reported by **[Chute et al. 1962] [Shafi. 1995]** which proved that the *Pasturella multocida* trivalent vaccine can gave high mean protection rate for controlling Fowl Cholera in Egypt.

Also protection rates of 83.3% and 93.3% were recorded in immunized chickens after challenge with virulent *Avibacterium paragallinarum* serotype A and C as shown in **table(2)**. These results agreed with those reported by **[Glisson. 1998] [Mouahid et al. 1991] [Nakamura et al. 1994]**, who found that chickens vaccinated with *Avibacterium paragallinarum* autogenous bacterin showed a very broad immune response and protection against *A. paragallinarum* infection.

Data represented in **table(3)**, showed The protection rate recorded in chickens immunized with the prepared vaccine against challenge with virulent *Ornithobacterium* rhinotracheale serotype A (96.6%) agreed with those reported by other authors **[Schuijffel et al. 2006] [Murthy et al. 2007]** who proved that vaccination of broiler at 6^th^ of age and followed by booster dose can effectively protect against *Ornithobacterium* rhinotracheale infection. It has been reported that natural protection against ORT infection is largely based on the development of humoral immune response **[Schuijffel et al. 2006]**.

Also, it is interesting that no re-isolation of *Pasturella multocida, Avibacterium paragallinarum* and *Ornithobacterium rhinotracheale* were recorded from respiratory and internal organs in vaccinated chickens. While the re-isolation of these bacterial strains was successful in control non vaccinated chickens.

Finally, It has been recorded in the present study that the use of trivalent vaccines against virulent serotypes of the following pathogens *Pasturella multocida, Avibacterium paragallinarum* and *Ornithobacterium rhinotracheale* combined with the use of strong adjuvant (Montanid ISA70) induced strong humoral immune responses and protective immunity in the vaccinated chickens. Thus this vaccine can be considered as a strong approach to the control of respiratory infections caused by these pathogens.

